# Cell type-specific Extracellular Vesicles in Mouse Brain: Proteomic Signatures Highlight Astrocytic GlialCAM Network and GPCR Enrichment

**DOI:** 10.64898/2025.12.12.693760

**Authors:** Alba M. Lucart-Sanchez, Edward Sellés-Climent, Jorge Navarro-Calvo, Guillem Pont-Espinós, Jose A. Gomez-Sanchez, Raúl Estévez, Luis M. Valor, Rocío Pérez-González

**Affiliations:** Instituto de Investigación Sanitaria y Biomédica de Alicante (ISABIAL), 03010, Alicante, Spain; Instituto de Neurociencias de Alicante, Universidad Miguel Hernández-CSIC, 03550, San Juan de Alicante, Spain; Centro de Investigación en Red-Enfermedades Neurodegenerativas (CIBERNED), Madrid, Spain; Physiology Unit, Department of Physiological Sciences, School of Medicine and Health Sciences, Institute of Neurosciences, University of Barcelona, 08907 L’Hospitalet de Llobregat, Spain; Neuroscience program, Physiology and pathology of the functional relationship between glia and neurons-IDIBELL, L’Hospitalet de Llobregat, Spain; The Spanish Center of Rare Diseases (CIBERER U-731), Baldiri Reixac 10, E-08028 Barcelona, Spain; Instituto de Investigación, Desarrollo e Innovación en Biotecnología Sanitaria de Elche (IDiBE), Elche, Spain

**Keywords:** extracellular vesicles, astrocytes, proteomics, GPCR, GlialCAM, blood-brain barrier

## Abstract

Extracellular vesicles (EVs) mediate intercellular communication in the central nervous system (CNS) and are emerging as biomarkers of brain health and disease. However, the molecular composition of cell type-specific brain EVs, particularly astrocyte-derived EVs (ADEVs), remains poorly defined. We performed comparative proteomic analysis of neuronal (NDEVs), microglial (MDEVs), and astrocytic (ADEVs) from mouse brain using magnetic immunocapture and LC-MS/MS proteomic profiling. Each EV subtype displayed distinct molecular fingerprints. NDEVs were enriched in synaptic and neurogenesis-related proteins (e.g., APP, SNAP25, GPR158 and BDNF), whereas MDEVs contained immune and phagocytic markers (e.g., TMEM119, CX3CR1, CD11B). Strikingly, the ADEV proteome closely mirrored the recently characterized GlialCAM interactome from leukodystrophy research, encompassing GlialCAM/MLC1 and associated partners involved in ion and water homeostasis (EAAT1/2, AQP4, GJA1), together with GPCRs such as GPRC5B. This overlap suggests that ADEVs encapsulate a molecular scaffold characteristic of astrocytic endfeet, potentially extending their signaling functions to the extracellular space. In conclusion, our study provides a detailed comparative proteomic characterization of brain cell type-specific EVs, revealing that ADEVs contain the GlialCAM/MLC1 network and GPCRs. These findings highlight ADEVs as promising candidates for biomarker development and mechanistic studies of blood-brain barrier integrity and neurodegenerative disorders.

## INTRODUCTION

Extracellular vesicles (EVs) are nanosized, lipid-bilayer particles secreted by all cell types in the central nervous system (CNS), including neurons, astrocytes, and microglia. They can carry proteins, lipids, and nucleic acids reflective of their cells of origin, thereby serving as dynamic mediators of intercellular communication and signaling ^1^. Within the brain, EVs play essential roles in maintaining CNS homeostasis and facilitating neurovascular and neuroimmune signaling ^2^. Importantly, EVs secreted by brain cells can cross the blood–brain barrier (BBB) bidirectionally, enabling molecular exchange between the CNS and peripheral circulation under both physiological and pathological conditions ^3–6^. This property positions brain EVs as promising, minimally invasive biomarkers of CNS diseases, neuroinflammation, and BBB function.

Despite this potential, isolating cell type–specific brain EVs from biofluids remains a major technical challenge. This is because EV surface proteins that enable antibody-based immunocapture must be both cell-specific and extracellularly accessible, criteria that are difficult to satisfy given the molecular overlap between CNS and peripheral EVs ^7^. Moreover, the intrinsic cellular heterogeneity of the brain, encompassing diverse neuronal and glial populations, further complicates efforts to achieve precise and reproducible identification of EV subtypes.

To date, neuronal-derived EVs (NDEVs) have primarily been isolated using L1CAM; however, the predominance of soluble L1CAM fragments over full-length forms has raised concerns regarding its specificity ^8,9^. Additional neuronal markers, such as ATP1A3, have shown promise for improving NDEVS isolation ^10^. For microglia-derived EVs (MDEVs), CD11b ^11,12^ and TMEM119 ^13,14^ have been used. In contrast, relatively few studies have systematically examined astrocyte-derived EVs (ADEVs), despite astrocytes’ central roles in neurovascular coupling, BBB integrity, and metabolic support of neurons. Markers such as EAAT1, EAAT2, and ATP1B2 have been used to enrich for ADEVs ^15–17^, with EAAT1 emerging as the most widely applied ^18^, while Aquaporin-4 (AQP4) has been proposed as a new immunocapture target ^15^. However, the surface proteome of astrocyte-derived EV isolated from brain tissue, particularly proteins associated with astrocytic endfeet and neurovascular interfaces, remains incompletely defined.

Here, we address this knowledge gap by performing a comparative proteomic analysis of three major brain EV subtypes: neuronal (NDEVs), microglial (MDEVs), and astrocytic (ADEVs), isolated from the same mouse brains using immunocapture and LC-MS/MS. Our results reveal distinct molecular signatures that reflect the specialized physiological roles of each parent cell type. Notably, ADEVs exhibit a proteomic profile that strongly overlaps with the GlialCAM interactome, including GlialCAM/MLC1, GPCRs, and transporters/ion channels, suggesting that ADEVs selectively encapsulate components of astrocytic endfeet function and signaling hubs. These findings provide a molecular framework for refining immunocapture strategies and advancing the use of ADEVs as biomarkers of BBB integrity and CNS pathology.

## MATERIALS AND METHODS

### Mouse housing and brain tissue collection

C57BL/6J young adults were housed under a 12-h light/dark cycle with food and water provided ad libitum at the animal facilities of Servicios Técnicos de Investigación, Universidad de Alicante. After sacrifice, mouse brains were immediately dissected and snap-frozen. All procedures were approved by the Comité de Ética de la Investigación, Universidad de Alicante, and authorized by the Dirección General de Producción Agrícola y Ganadera, Generalitat Valenciana, according to European, national and regional laws.

### Extracellular Vesicle Isolation

Frozen mouse brains, without the cerebellum and olfactory bulbs, from female and male young adults, were used for EV isolation and characterization. Small EVs were isolated as previously described ^19,20^ with some modifications. Briefly, the tissues were minced and incubated with 20 U/mL papain (Worthington Biochemical Corporation, LK003176) and 40 U/ml DNAse I (Roche, 11284932001) in Hibernate A medium (ThermoFisher Scientific, A12475-01) for 15 min at 37°C. The enzymatic digestion was stopped by the addition of ice-cold Hibernate A supplemented with a cocktail of protease and inhibitors (5 μg/ml leupeptin, 5 μg/ml antipain dihydrochloride, 5 μg/ml pepstatin A, 1 mM phenylmethylsulfonyl fluoride (PMSF), 1 μM E-64, all from Sigma-Aldrich).

The solution was gently dissociated by pipetting and centrifuged at 300 x *g* for 10 min at 4°C to pellet undigested tissue and intact cells. The supernatant was subsequently filtered through a 40 μm cell strainer and then centrifuged at 2,000 x *g* for 10 min at 4°C to discard large debris and apoptotic bodies. The resulting supernatant was centrifuged at 10,000 x *g* for 30 min at 4°C to discard smaller debris, passed through a 0.2 μm surfactant-free cellulose acetate (SFCA) membrane filter (Corning, 431219), and ultra-centrifuged at 100,000 x *g* for 70 min at 4°C to pellet EVs in a Beckman Coulter 70 Ti fixed-angle rotor (k factor 106). The pellet was washed in phosphate-buffered saline (PBS), re-centrifuged at 100,000 x *g* for 70 min at 4°C and resuspended in PBS for subsequent immunocapture steps.

### Immunocapture of Cell Type–Specific EVs (ATPA3⁺, CD11b⁺, ACSA-2⁺)

Magnetic beads (Invitrogen, 14311D) were conjugated with anti-ATP1A3 (Invitrogen, cat. no. MA3-915) and anti-CD11b (BioLegend, 101201) for neuronal-derived EVs (NDEVs) and microglial-derived EVs (MDEVs), respectively, at a ratio of 5 µg of antibody per mg of beads. Astrocyte-derived EVs (ADEVs) were immunocaptured using the anti-ACSA-2 Microbead Kit (Miltenyi Biotec, 130-097-679). Brain EVs (45 µg) were incubated with 10 µL of FcR Blocking Reagent, Mouse (Miltenyi Biotec, 130-092-575) for 15 min on ice. Subsequently, 100 µL of antibody-conjugated magnetic beads were added for overnight incubation at 4 °C under rotation. Bulk EVs and unbound fractions corresponding to the non-immunocaptured EVs were kept for assessing the immunocapture efficiency, while beads incubated with only PBS (1×) were used as negative controls. For electron microscopy, nanoparticle tracking analysis (NTA), and mass spectrometry, cell type–specific EVs were released from magnetic beads using IgG Elution Buffer (Thermo Fisher Scientific, 21028), followed by pH neutralization with 1 M Tris base buffer.

### Sample lysis, protein estimation and Western Blotting

Tissue pellets and EVs were lysed in 2× RIPA buffer (except for immunocaptured EVs that were eluted in 1× RIPA) supplemented with protease inhibitors and sonicated for 45 sec in a water bath at 4 °C followed by incubation on ice for 20 min with brief vortexing every 2–3 min. The first pellets obtained after centrifugation at 300 × g were mechanically dissociated using a pestle on ice before sonication, centrifuged at 15,700 × g for 15 min at 4 °C, and the supernatant lysed for protein analysis. Protein concentrations were determined using the Pierce BCA Protein Estimation Assay Kit (Thermo Scientific, 23227). Lysates were then denatured in 6X Laemmli SDS sample buffer (Thermo Scientific, J61337-AC), under reducing condition, and proteins were separated by SDS–PAGE using 10% TGX Stain-Free Fast Cast acrylamide gels (Bio-Rad, catalog no. 1610183). Magnetic beads (except microbeads) were removed before SDS-PAGE. Proteins were transferred onto 0.45 µm PVDF membranes using the Trans-Blot Turbo Mini-Size LF PVDF Transfer Pack (Bio-Rad, 10026934). Membranes were blocked with EveryBlot Blocking Buffer (Bio-Rad, 12010020) and incubated overnight at 4 °C with the following primary antibodies: anti-ATP1A3 (Rabbit, 1:1000, Proteintech, 10868-1-AP), anti-Glutamine Synthetase (Rabbit, 1:1000, GeneTex, GTX109121), anti-Alix (Rabbit, 1:1000, Cell Signaling Technology, 92880S), anti-CD11b (Rabbit, 1:1000, GeneTex, GTX640543), anti-Calnexin (Rabbit, 1:20.000, GeneTex, GTX109669). For the analysis of the astrocytic GlialCAM interactome proteins, we used anti-GlialCAM ^21^, anti-GPCR5B ^22^ and anti- MLC1 ^23^, all Rabbit at 1:500 dilution. After washing, membranes were incubated for 1 h at room temperature with horseradish peroxidase (HRP)-conjugated secondary antibody (anti-rabbit IgG, Cell Signaling Technology, 7074S). Protein bands were visualized using SuperSignal West Femto Maximum Sensitivity Substrate (Thermo Scientific, A38554) and imaged with the ChemiDoc MP Imaging System (Bio-Rad). Densitometric analysis was performed using ImageJ software (NIH).

### Scanning Transmission Electron Microscopy (STEM) of EV Preparations

EV imaging was performed at the Microscopy Unit of ISABIAL. For STEM sample preparation, 10 µL of EV suspension was placed onto Formvar/Carbon-coated copper grids (200 mesh) and allowed to adsorb for 25 min. Excess liquid was gently removed with filter paper. To enhance surface contrast, the samples were negatively stained with 10 µL of 2% uranyl acetate (pH 7.4) for 20 s. The excess stain was carefully removed without directly contacting the grid surface. After complete air-drying, the grids were mounted in a dedicated STEM grid holder and examined using a field emission scanning electron microscope (FE-SEM, Zeiss GeminiSEM 460) equipped with a STEM detector operating at 30.0 kV. Images were acquired at magnifications of 10 µm and 3 µm to assess EV morphology and surface characteristics.

### Nanoparticle Tracking Analysis (NTA)

The particle size distribution and concentration measurements were evaluated with a Nanosight NS300 (Particle Tracking Analysis) instrument (Malvern Panalytical, Malvern, UK). EVs were resuspended in PBS and diluted to the working range of the system (106-109 particles/ml). Videos were captured and analysed with the Nanosight NS300 software (version 3.4) using a sCMOS camera.

### Mass spectrometry and proteomic data analysis

Mass spectrometry and proteomic analysis were performed in the proteomics facility of SCSIE University of Valencia, a member of Proteored. All protein extracts were concentrated to 20 µL using a SpeedVac and subsequently mixed with 25 µL of 4× Laemmli sample buffer (Bio-Rad, 1610747). Samples were vortexed for 5 min and heated at 95 °C for 5 min to ensure complete denaturation. The resulting protein extracts were loaded onto SDS–PAGE gels to assess sample quality, visualize protein bands, and estimate relative protein concentration.

Tandem mass spectrometry (LC–MS/MS) analysis was performed using a NanoElute2 HPLC system (Bruker) coupled to a timsTOF SCP mass spectrometer (Bruker). A total of 50 ng of digested peptides were separated on an Aurora Ultimate C18 analytical column (25 cm × 75 µm, 1.7 µm; Bruker Daltonics) using a 66-min linear gradient on the NanoElute2 system. Peptides were ionized via CaptiveSpray at 1700 V and 200 °C and analyzed in data-dependent acquisition mode (ddaPASEF) using the method SCP_dda_low_SampleAmount_5PASEF. Instrument parameters included a 1/K₀ range of 0.7–1.3 V·s/cm², ramp time of 166 ms, duty cycle of 100%, ramp rate of 5.78 Hz, MS averaging of 1, and auto-calibration disabled. The MS scan range was set to 100–1700 m/z with positive ion polarity and PASEF scan mode. MS/MS acquisition included five PASEF ramps per cycle with a total cycle time of 1.04 s, charge range of 0–5, target intensity of 20,000, an intensity threshold of 500, and active exclusion enabled.

Raw data files were analyzed using FragPipe to identify proteins and perform label-free quantification (LFQ) using standard in-house parameters. The combined_protein.tsv output file was used for statistical analysis. Contaminant, species-mismatched proteins, and those not consistently identified or quantified within the same condition were excluded. MaxLFQ intensity values were log₂-transformed, and missing values were imputed using the Missing Not At Random (MNAR) approach, which samples random values from a left-shifted Gaussian distribution (1.8 standard deviations downshift, width = 0.3). Differential expression analysis was performed using protein-wise linear models combined with empirical Bayes statistics implemented in the *limma* package (R/Bioconductor). Proteins with an adjusted *p*-value < 0.05 (FDR correction) and an absolute log₂ fold change ≥ 1 were considered significantly differentially enriched.

Venn diagrams were generated using InteractiVenn ^24^, gene ontology (GO) enrichment analysis was performed with ShinyGO 0.82 ^25^, and protein–protein interaction (PPI) networks were constructed using STRING v12.0 ^26^. The heatmaps were generated using GraphPad Prism (version 8.01; GraphPad Software, San Diego, CA, USA) except for Figure 3G-H, which was made with Excel. The volcano plots were generated using SRplot ^27^.

The mass spectrometry proteomics data have been deposited to the ProteomeXchange Consortium via the PRIDE ^28^ partner repository with the dataset identifier PXD071055.

### Meta-analysis with gene expression datasets

Differential enriched proteins were screened for cell-specific markers that derived from a curated list of 100-gene markers for neurons, astrocytes, oligodendrocytes, endothelial cells and microglia that are shared between humans and mice and that have been further validated in the literature (Supplemental File 2 of the original publication ^29^). To infer the cellular origin of the VE-derived proteins, we used the median gene expression from single cell transcriptomics of mouse brains (cortex and hippocampus) for the Allen Brain Atlas (https://brain-map.org/our-research/cell-types-taxonomies/cell-types-database-rna-seq-data/mouse-whole-cortex-and-hippocampus-10x), according to the last cellular taxonomy ^30^. Genes with median expression = 0 were removed, assuming an issue regarding sequencing coverage.

### Statistical Analysis

All statistical analyses were conducted using GraphPad Prism software (version 8.01; GraphPad Software, San Diego, CA, USA). A p-value < 0.05 was considered statistically significant.

## RESULTS

### Cell type-specific EVs Immunocapture Validation

To characterize cell type-specific small EVs from brain tissue, we optimized a workflow to enrich in neuronal, microglial, and astrocytic EVs, hereafter referred to as NDEVs, MDEVs and ADEVs, respectively, following Minimal information for studies of extracellular vesicles (MISEV2023) guidelines ^31^ and recommendation for EV studies from solid tissue ^32^. EV isolation was performed by differential ultracentrifugation followed by immunocapture of specific EVs (Supplementary Fig. 1). Throughout the isolation process, intermediate pellets were kept and analyzed by Western blot to assess the purity of the 100,000 x *g* pellet corresponding to bulk EVs. Bulk EVs were enriched in canonical EV markers Alix and CD63 (Supplementary Fig. 2A-B). Calnexin, an intracellular marker, was mostly excluded from bulk EVs, confirming minimal intracellular contamination (Supplementary Fig. 2A-B).

Western blot analysis verified the enrichment of ATP1A3, CD11b and glutamine synthetase in NDEVs, MDEVs and ADEVs, respectively (Fig. 1A-D). Nanoparticle tracking analysis (NTA) revealed distinct size and number distributions among EV populations (Fig. 1E-H). All EV subtypes exhibited a predominant size below 200 nm, with MDEVs exhibiting a slightly larger modal size at 181 ± 20.12 nm (Fig. 1G). In terms of EV numbers, ADEVs displayed the highest particle concentration of 8.87 × 10L particles/mL, followed by NDEVs (1.32× 10L particles/mL) and MDEVs (8.43 × 10^8^ particles/mL). Lastly, electron microscopy imaging corroborated the presence of intact vesicles with morphology and size consistent with small EVs in all populations (Fig. 1I).

**Figure 1.**
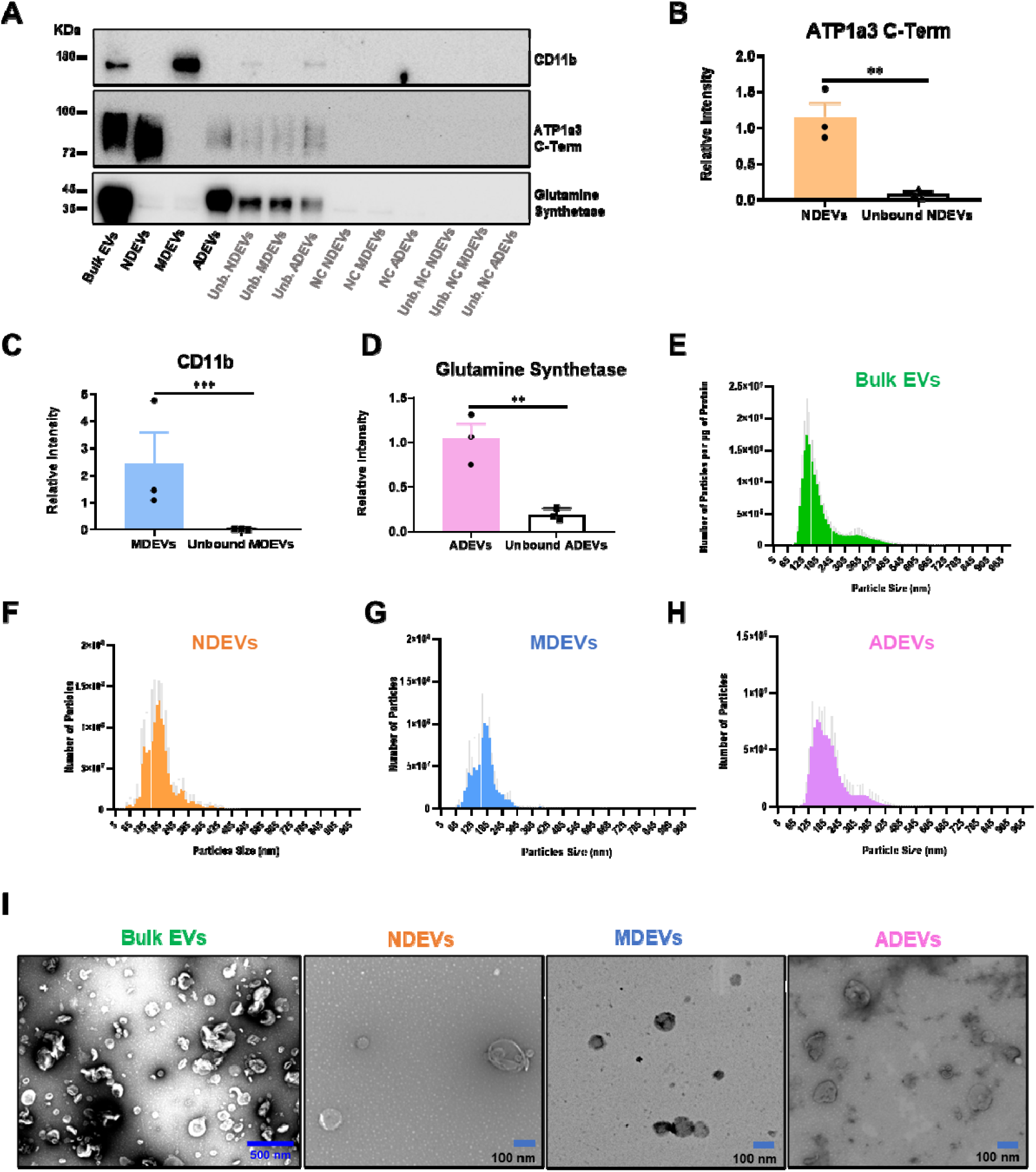
**Cell type-specific EVs Immunocapture Validation**. **A.** Representative Western blot showing enrichment of cell-specific markers: CD11b in MDEVs (CD11b_⁺_EVs), in NDEVs and glutamine synthetase in ADEVs (ACSA-2_⁺_EVs), compared to bulk EVs. Unbound fractions are also shown. Beads conjugated with the respective antibodies and incubated with only PBS (1×) were used as negative controls. **B.** Quantification of the relative intensity (mean ± SD) of immunocaptured EVs (bound) versus unbound fractions confirmed the efficiency of the immunocapture. Student t-test, N=3, ** p<0.01, *** p < 0.001. **C.** Size and number distributions among the cell type–specific EV populations by Nanoparticle tracking analysis (NTA) (N=4 per cell type-specific EV), the data graphics represented the mean of the number of particles per Bin center (color bars) and the mean ± SEM (standard error of the mean; light gray bars). The number of particles in bulk EVs was normalized by total _μ_g of EV protein estimated by brain weight. **D.** Scanning transmission electron microscopy (STEM) images show vesicles with size and morpholog consistent with EVs for each cell type–specific preparation. NDEVs – Neuron-derived EVs; MDEVs – Microglia-derived EVs; ADEVs – Astrocyte-derived EVs.

### Protein profiling of bulk, neuronal, microglial, and astrocytic EVs

Next, we conducted a comparative proteomic analysis of the three cell type-specific populations together with bulk EVs. Principal component analysis (PCA) revealed distinct sample distributions among EV subtypes, with segregated clustering patterns and high reproducibility across biological replicates (Fig. 2A). A heatmap of log_2_-centered protein intensities further illustrated different protein expression profiles among NDEVs, MDEVs, ADEVs and bulk EVs, with high similarity in protein profiles observed within replicates (Fig. 2B).

**Figure 2.**
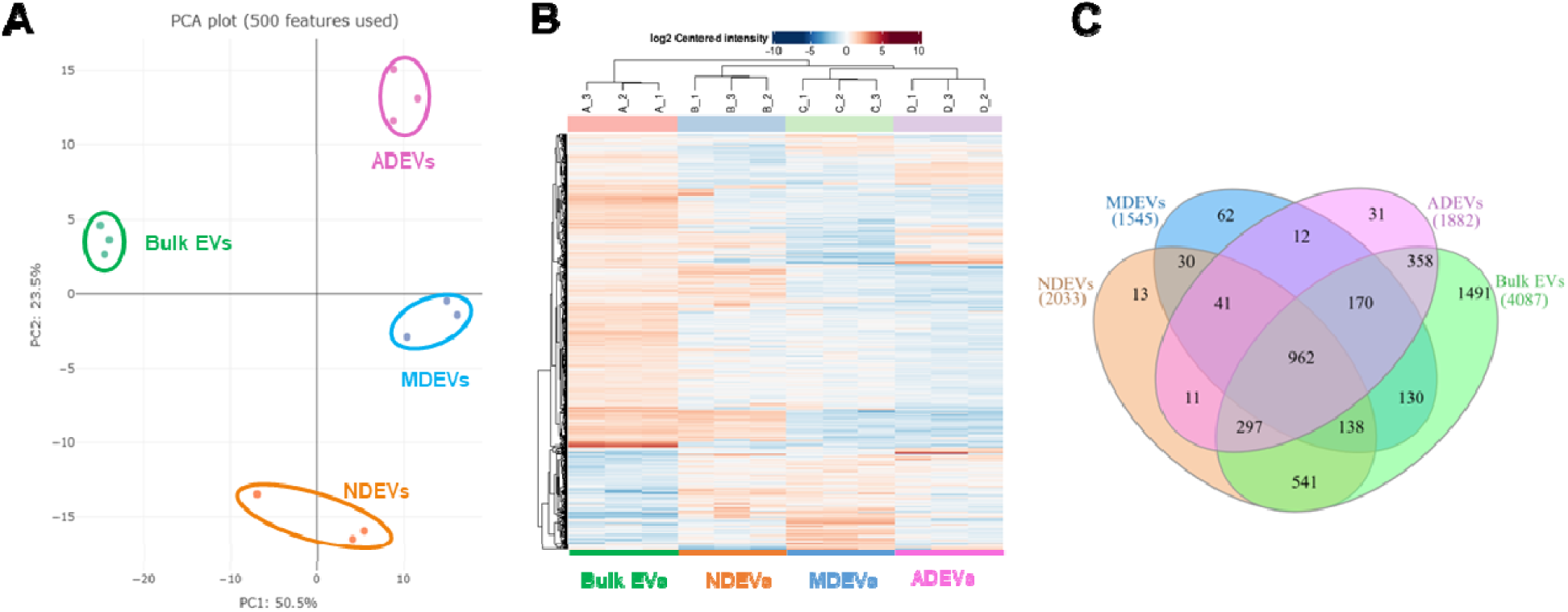
Measurement of proteins that provide separation among cell type-specific and bulk EVs in the brain. **A.** Principal component analysis (PCA) based on the top 500 most variable proteins reveals clear clustering by EV subpopulation group, indicating distinct proteomic profiles among NDEVs, MDEVs, ADEVs, and bulk EVs. **B.** Heatmap of log2-centered protein intensity values showing distinct expression patterns specific to each cell type-specific EVs, and high reproducibility across biological replicates (N = 3 per group). **C.** Venn diagram illustrating the distribution of proteins identified across NDEVs, MDEVs, ADEVs, and Bulk EVs. This diagram was performed based on MaxLFQ Intensity.

**Figure 3.**
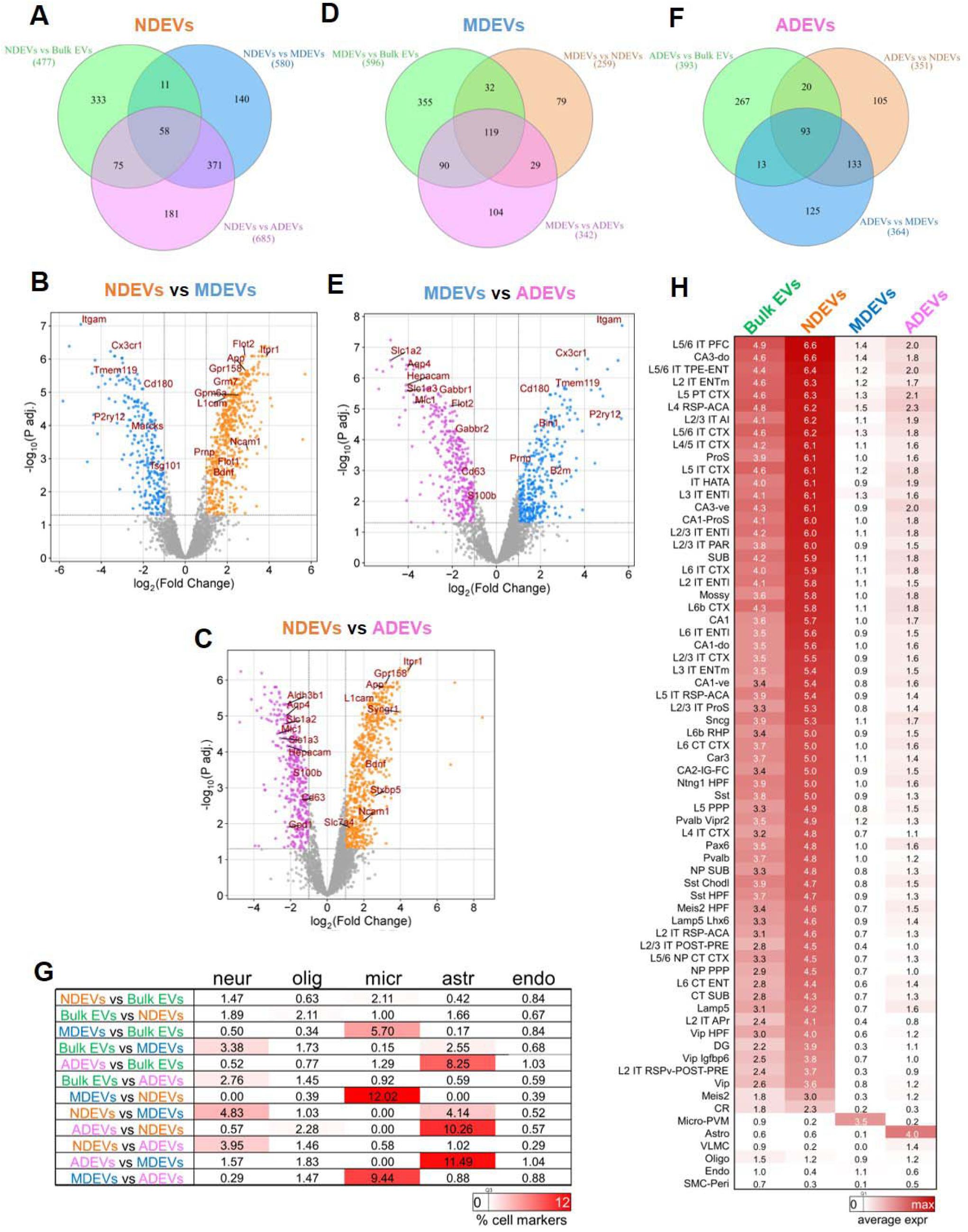
Differential Protein Signatures Across Cell type-specific EVs and Bulk EVs. **A.** Venn diagram illustrating proteins upregulated in NDEVs relative to bulk EVs and other cell type-specific EV subtypes, revealing 58 uniquely upregulated proteins. **B.** Volcano Plot comparing the upregulated proteins in NDEVs compared to MDEVs, showing the upregulated proteins in NDEVs versus MDEVs (GPR158, GRM7, ITPR1, APP, GPM6A, L1CAM, NCAM1, PRNP, FLOT1, and BDNF) and upregulated in MDEVs compared to NDEVs (CD11b (Itgam), CX3CR1, TMEM119, CD180, P2RY12, MARCKs, and TSG101). **C.** Volcano Plot comparing the upregulated proteins in NDEVs compared to ADEVs, showing the upregulated proteins in NDEVs versus ADEVs (GPR158, ITPR1, APP, L1CAM, SYNGR1, BDNF, STXBP5, NCAM1, and SLC7A4) and upregulated in ADEVs compared to NDEVs (ALDH3B1, AQP4, SLC1A2, MLC1, SLC1A3, HEPACAM, S100B, CD63, and GPD1). Note: GRM7 was also upregulated in NDEVs relative to ADEVs but it is not shown because its parameters were nearly identical to APP, causing an overlap. **D.** Venn diagram showing proteins upregulated in MDEVs compared with the remaining EV subtypes and bulk EVs, identifying 119 upregulated proteins. **E.** Volcano Plot comparing the upregulated proteins in MDEVs compared to ADEVs, showing the upregulated proteins in MDEVs versus ADEVs (CD11b (Itgam), CX3CR1, TMEM119, CD180, P2RY12, BIN2, PRNP, and B2M) and upregulated in ADEVs compared to MDEVs (SLC1A2, AQP4, HEPACAM, SLC1A3, MLC1, FLOT2, GABBR1-2, CD63, and S100B). **F.** Venn diagram depicting the proteins upregulated in ADEVs, with 93 proteins elevated in this cell type-specific EV population. **G.** Heatmap displaying the percentage of brain cell-specific markers described from a transcriptomics meta-analysis. neur = neuron; olig = oligodendrocyte; micr = microglia; astr = astrocyte; endo = endothelial cells. **H.** Heatmap showing the average expression of upregulated NDEVs, MDEVs or ADEVs proteins in different cell-type clusters defined by scRNA-seq (Allen brain atlas). Volcano plots show gene names instead of protein names.

After applying stringent filtering criteria (see Material and Methods), a total of 4287 proteins were evaluated. Most of the proteins were identified in bulk EVs (4087), while 2033 were detected in NDEVs, 1545 in MDEVs, and 1882 in ADEVs. A core set of 962 proteins, including canonical EV proteins such as ALIX, CD9, CD81, members of the Annexin family, and FLOT1, were shared across all EV groups (Fig. 2C). Exclusive proteins accounted for 1491 entries in bulk EVs, representing approximately 35% of all proteins. Among cell type-specific EVs, 13 proteins were unique to NDEVs (e.g., VAMP7 and SLC7A4), 62 to MDEVs (e.g., GPR34, CLEC5A, and CLEC4A2), and 31 to ADEVs (e.g., LPR4 and KCNJ16). The complete list of proteins can be found in Supplementary Table 1.Overlap analysis revealed that NDEVs shared the highest number of proteins with bulk EVs (541), including AMPA receptor subunits GRIA1, GRIA2, and GRIA3, whereas MDEVs displayed the most distinct profile, sharing only 130 proteins (e.g., CX3CR1 and TMEM119) (Fig. 2C). These findings indicate that bulk EVs are predominantly neuronal in origin, while microglial EVs are the most underrepresented population.

### Differential protein expression profile in cell type-specific EVs

To further characterize the EV subtypes, differential expression analyses were performed among bulk EVs and cell-type–specific EVs using thresholds of Log2 fold-change ≥ 1 and an adjusted p-value < 0.05.

NDEVs exhibited 58 proteins exclusively upregulated compared to all other groups, including BDNF, GRM7, ITPR1, and GPR158 (Figure 3A-C, Supplementary Table 2). Gene Ontology (GO) Biological Process enrichment analysis revealed that these proteins were primarily involved in localization, transport, transmembrane and ion transport, and cell–cell signaling (Supplementary Fig. 3). In pairwise comparisons, NDEVs showed 371 proteins upregulated relative to MDEVs and ADEVs, including the well-known NDEV markers NCAM1 and L1CAM, as well as neurodegeneration-associated proteins such as APP (Fig. 3A-C). Additionally, 181 were elevated relative to ADEVs (e.g., SYNGR1 and STXBP5; Fig. 3A, C) and 140 proteins versus MDEVs (e.g., GPM6A, FLOT1; Fig. 3A, B). Notably, 11 proteins were uniquely upregulated in NDEVs compared with both bulk EVs and MDEVs (e.g., FLOT2; Fig. 3A, B), and 75 proteins were shared as upregulated relative to bulk EVs and ADEVs (e.g., SLC7A4; Fig. 3A, C; Supplementary Table 2).

MDEVs contained 119 proteins upregulated compared to bulk EVs and other cell type-specific EVs, which included the microglial markers ITGAM, CX3CR1, TMEM119, and P2RY12 (Fig. 3D, B, E). These proteins were enriched in pathways related to cell activation, immune processes, and vesicle-mediated transport (Supplementary Fig 3). Pairwise comparisons identified 79 proteins elevated versus NDEVs (e.g., TSG101 and MARCKS; Fig. 3D, B) and 104 versus ADEVs (e.g., PRNP, B2M, BIN1; Fig. 3D, E), with 29 proteins consistently increased in MDEVs compared with both NDEVs and ADEVs (e.g., CD180; Fig. 3D, C, E Supplementary Table 3).

ADEVs displayed 93 proteins upregulated relative to bulk EVs, NDEVs and MDEVs, including astroglial markers AQP4, S100B, EAAT2, and GlialCAM (HEPACAM) (Fig. 3F, E, C). GO analysis indicated enrichment for amino acid and organic acid transport, as well as chemical homeostasis (Supplementary Fig. 3). Pairwise comparisons, 133 proteins were upregulated versus NDEVs and MDEVs, including EAAT1, CD63, and MLC1 (Fig. 3F, E, C). Specifically, 125 proteins were elevated versus MDEVs (e.g., GABBR1-2 and FLOT2; Fig. 3F, E) and 105 versus NDEVs (e.g., GPD1-2, ALDH3B1; Fig. 3F, C; Supplementary Table 4).

To further validate cell type-specificity of the retrieved EV proteins, we examined the presence of curated top 100 cell-specific expressed genes that were described for each major brain cell type ^29^ across the pairwise comparisons. ADEVs and MDEVs showed strong overlap with reported markers, whereas NDEVs enrichment was more moderate (Fig. 3G), consistent with their similarity to bulk EVs. In contrast, integration with single-cell RNAseq data from the Allen Brain Atlas (see Material and Methods), confirmed representation of major neuronal-cell types and brain regions in our NDEVs proteome (Fig.3H), i.e., no apparent enrichment for specific neuronal subtypes were observed; this analysis also supported the specificity of MDEV and ADEV profiles (Fig.3H). Notably, differentially expressed proteins in bulk EVs were mainly of neuronal origin, although their averaged gene expression was lower compared to NDEV corresponding genes, with poor representation of oligodendrocyte, endothelial, and pericyte gene markers (Fig. 3H).

### Distribution of the commonly identified EV markers across cell type-specific EVs

To characterize the distribution of known canonical EV markers among the different EV populations, we compared our detected proteins with the ExoCarta Top 100 small EV protein list (http://exocarta.org/sEV_top100). Of the 100 catalogued proteins, 83 were identified in all our datasets. Common markers included Annexins (ANXA1, ANXA2, ANXA5, ANXA6), tetraspanins (CD9, CD81), and the Endosomal Sorting Complex Required for Transport (ESCRT)-accessory protein ALIX. Three proteins (FLOT2, TFRC, MYO1C) were shared between NDEVs and ADEVs, three (RDX, TSG101, EHD1) overlapped between MDEVs and ADEVs, and one (CD63) was detected only in ADEVs (Fig. 4A; Supplementary Table 5).

**Figure 4.**
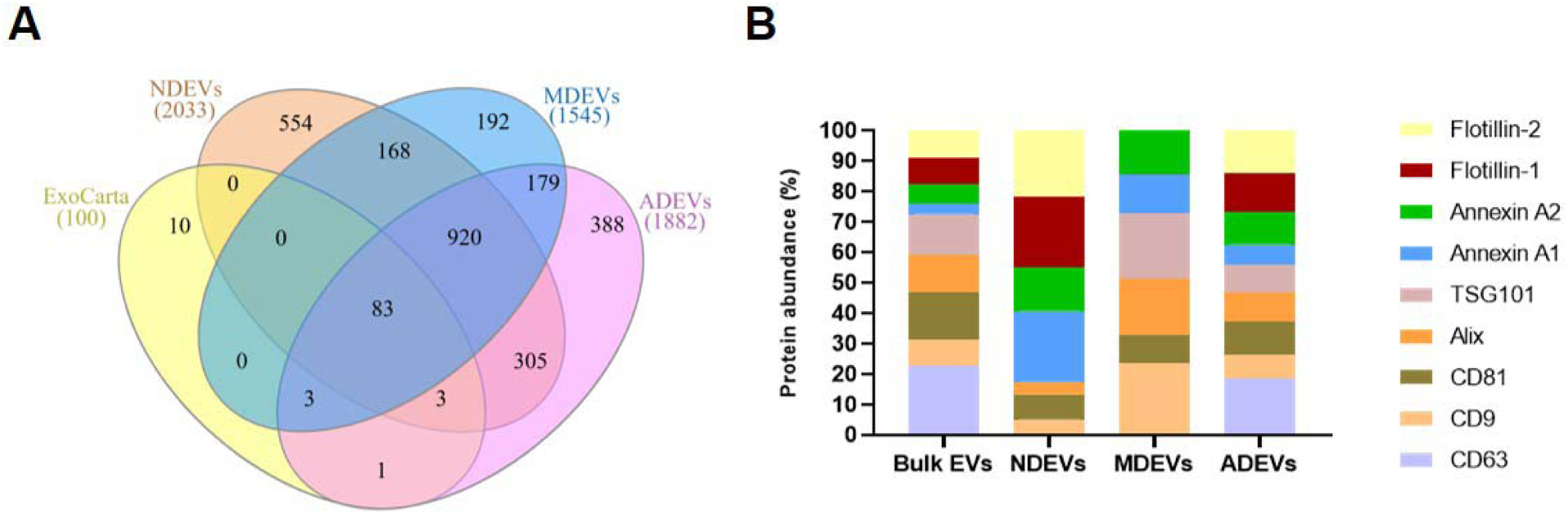
Distribution of Commonly Identified EV Markers Across Cell type-specific EVs. **A.** Venn diagram showing the overlap between proteins identified in NDEVs, ADEVs, MDEVs, and the ExoCarta Top 100 small EV protein list. A total of 83 proteins were shared between the 3 cell-types–specific EV datasets and the ExoCarta reference list. **B.** Stacked bar chart showing the protein abundance (%) of common EV markers across cell-type–specific EVs and bulk EVs.

Flotillin family members (FLOT1, FLOT2) were significantly enriched in NDEVs (*p* adj ≤ 1.86 × 10⁻^3^; 8.14 × 10⁻^6^) and ADEVs (*p* adj ≤ 9.96 × 10⁻^3^; 8.06 × 10⁻^6^) compared to MDEVs. NDEVs also showed higher FLOT2 levels relative to bulk EVs (*p* adj = 6.18 × 10⁻L). Among Annexins, ANXA1 was elevated in NDEVs but not significantly different from ADEVs or MDEVs, while it was significantly enriched in bulk EVs compared to NDEVs and MDEVs (*p* adj = 1.09 × 10⁻² and 2.05 × 10⁻²). ANXA2 was consistently expressed across EV subtypes but significantly higher in all cell type–specific EVs compared to bulk EVs (vs NDEVs p adj = 7,36E × 10L^4^; vs MDEVs p adj = 2,16 × 10L^3^; vs ADEVs p adj = 0,0191).

Regarding tetraspanins, CD9 was markedly enriched in MDEVs relative to NDEVs (*p* adj = 2.66 × 10⁻L) and ADEVs (*p* adj = 9.89 × 10⁻L), as well as bulk EVs (*p* adj = 1.65 × 10⁻³). CD63 was only detected in ADEVs and bulk EVs, with no significant difference between them (*p* adj = 0.234). CD81 expression remained relatively stable but was higher in bulk EVs than in cell type–specific EVs (bulk EVs vs NDEVs p adj = 3,90E × 10L^3^; vs MDEVs p adj = 7,92 × 10L^4^; vs ADEVs p adj = 0,0141).

Finally, ESCRT-related proteins ALIX and TSG101, MDEVs were significantly enriched in MDEVs compared to NDEVs (*p* adj = 1.61 × 10⁻L and 3.19 × 10⁻L, respectively), with ALIX also elevated in ADEVs relative to NDEVs (*p* adj = 2.34 × 10⁻²). In contrast, MDEVs showed no significant enrichment relative to bulk EVs for these proteins. Table 1 summarizes fold-changes and *p* adj values for the expression of common EV markers across Cell type-specific EVs. The relative abundance of each EV marker across EV populations is shown in Figure 6B.

**Table 1.** Enrichment Patterns of Common EV Markers in Bulk and Cell type-specific EVs. . A. Enrichment of common EV markers across NDEVs, MDEVs, and ADEVs. Log2 fold changes (FC) and adjusted p-values (p-adj) are shown.

| A. - Common EV markers distributions across Cell type-specific EVs |  |  |  |  |  |  |  |
| --- | --- | --- | --- | --- | --- | --- | --- |
| Protein name | Gene name | log<sub>2</sub> FC NDEVs vs MDEVs | log<sub>2</sub> FC NDEVs vs ADEVs | log<sub>2</sub> FC MDEVs vs ADEVs | p adj NDEVs vs MDEVs | p adj NDEVs vs ADEVs | p adj MDEVs vs ADEVs |
| ANXA1 | Anxa 1 | 1,17 | 0,346 | 0,825 | 0,127 | 0,756 | 0,303 |
| ANXA2 | Anxa2 | -0,02 | 0,282 | 0,302 | 0,971 | 0,33 | 0,316 |
| CD63 | Cd63 | 0,175 | -1,44 | -1,62 | 7,74E-01 | 2,18E-03 | 1,70E-03 |
| CD81 | Cd81 | -0,128 | -0,533 | -0,405 | 0,752 | 0,0609 | 0,17 |
| CD9 | Cd9 | -2,24 | -0,77 | 1,47 | 2,66E-05 | 0,0343 | 9,89E-04 |
| FLOT1 | Flot1 | 2,62 | 0,486 | -2,13 | 0,51 | 0,118 | 9,96E-03 |
| FLOT2 | Flot2 | 2,84 | 0,758 | -2,08 | 8,14E-07 | 6,36E-03 | 8,06E-06 |
| ALIX | Pdcd6ip | -2,29 | -1,41 | 0,879 | 0,00129 | 0,0234 | 0,164 |
| TSG101 | Tsg101 | -1,76 | -0,813 | 0,946 | 0,00234 | 0,103 | 0,0772 |
| B.- Common EV markers distributions between Bulk EVs and Cell- type specific EVs |  |  |  |  |  |  |  |
| Protein name | Gene name | log<sub>2</sub> FC Bulk EVs vs NDEVs | log<sub>2</sub> FC Bulk EVs vs MDEVs | log<sub>2</sub> FC Bulk EVs vs ADEVs | p adj Bulk EVs vs NDEVs | p adj Bulk EVs vs MDEVs | p adj Bulk EVs vs ADEVs |
| ANXA1 | Anxa 1 | -2,01 | -1,66 | -0,837 | 1,09E-02 | 2,05E-02 | 0,204 |
| ANXA2 | Anxa2 | -0,916 | -0,936 | -0,634 | 3,90E-03 | 2,16E-03 | 0,0191 |
| CD63 | Cd63 | 1,86 | 2,04 | 0,42 | 5,11E-04 | 1,27E-04 | 0,234 |
| CD81 | Cd81 | 1,19 | 1,06 | 0,653 | 7,36E-04 | 7,92E-04 | 0,0141 |
| CD9 | Cd9 | 1,05 | -1,18 | 0,286 | 5,25E-03 | 1,65E-03 | 0,338 |
| FLOT1 | Flot1 | -1,03 | 1,59 | -0,541 | 0,118 | 1,81E-02 | 0,379 |
| FLOT2 | Flot2 | -1,11 | 1,73 | -0,355 | 6,18E-04 | 1,21E-05 | 0,11 |
| ALIX | Pdcd6ip | 1,93 | -0,363 | 0,516 | 3,56E-03 | 0,475 | 0,31 |
| TSG101 | Tsg101 | 1,36 | -0,395 | 0,551 | 8,61E-03 | 0,351 | 0,198 |

### G protein-coupled receptors (GPCRs) in Cell type-specific EVs

G protein-coupled receptors (GPCRs) control neurotransmission, synaptic plasticity, neuroinflammation, and neuromodulation, and their presence in EVs suggests that cells may export key signaling receptors to influence their environment ^33,34^. Our proteomic analysis revealed a broad representation of GPCRs across all EV subtypes, with distinct enrichment patterns specific to each EV subtype. We compared upregulated proteins (FC≥ 1 and p adj. <0.05) in the cell type-specific EVs and bulk EVs against the curated GPCR lists from the IUPHAR/BPS Guide to PHARMACOLOGY (https://www.guidetopharmacology.org/GRAC/ReceptorFamiliesForward?type=GPCR).

NDEVs showed prominent enrichment in Class C receptors, particularly metabotropic glutamate receptors (GRM1, GRM2, GRM5, GRM7, and GRM8) and the orphan receptor GPR158 (Fig. 5A-B). Further comparison of NDEVS GPCRs with bulk EVs identified GPR158 as the most significantly enriched GPCR in NDEVs. Its enrichment was statistically significant compared to bulk EVs (*p* adj. = 2.14 × 10⁻³), MDEVs (*p* adj.= 2.72 × 10⁻L), and ADEVs (*p* adj. = 1.27 × 10⁻L) (Fig. 5D-E). Other GPCRs enriched in NDEVs included the Adhesion Class receptors ADGRL1 and ADGRL3, the somatostatin receptor SS2R, and the cannabinoid receptor CNR1 (Fig. 5A-B). Specifically, GRM7 and ADGRL3 were also upregulated compared to bulk EVs (Fig. 5D-E).

**Figure 5.**
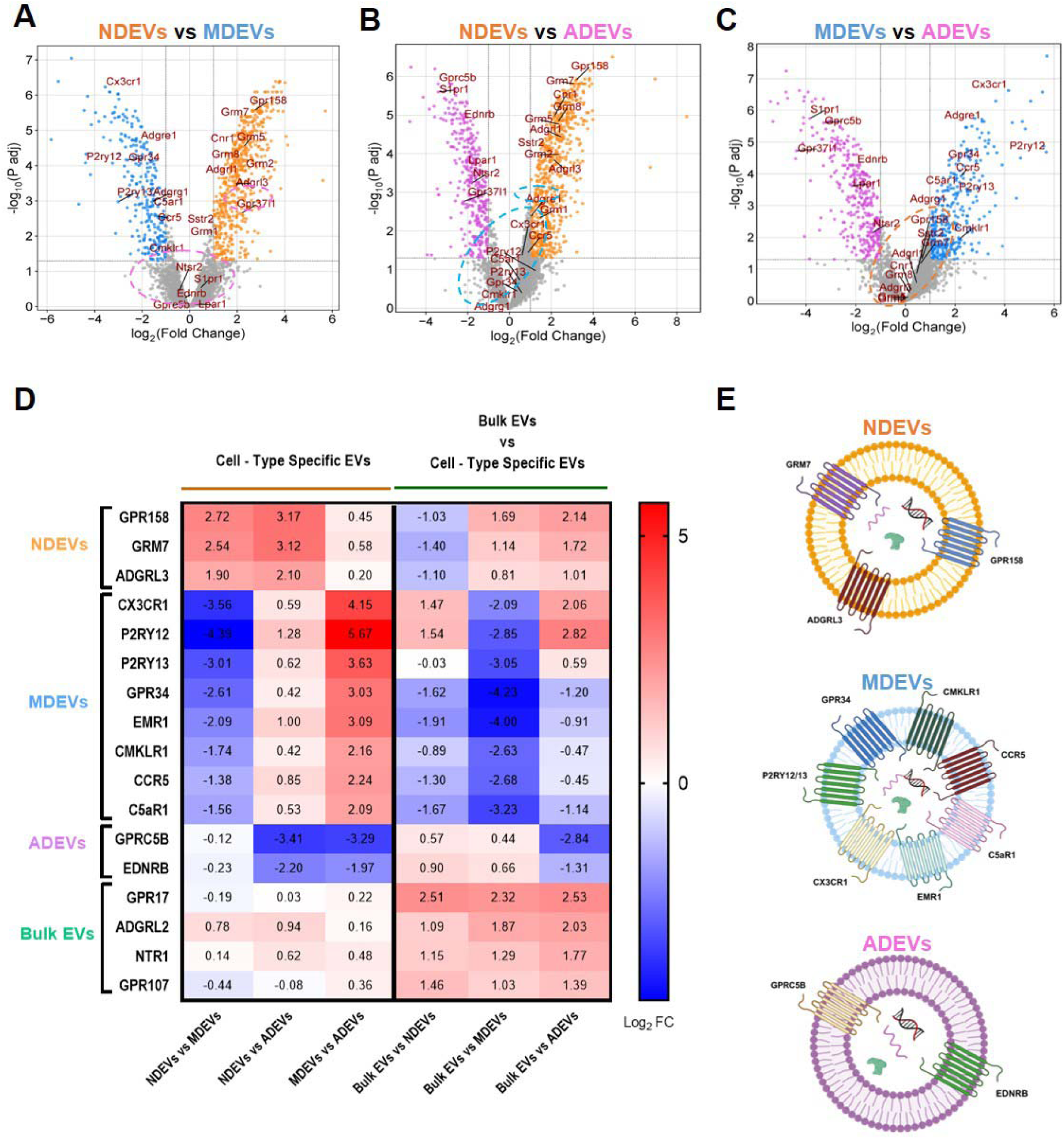
Differential Enrichment of G Protein-Coupled Receptors (GPCRs) in Cell type-specific EVs and Bulk EVs. A-C. Comparative Volcano Plots of Cell type-specific EV GPCR Enrichment. Volcano plots compare the relative abundance (Fold Change, Log2 FC) and statistical significance (p adj.) of upregulated GPCRs (FC≥ 1 and p adj. <0.05) between specific EV subtypes. **A.** NDEV vs MDEV: Shows GPCRs significantly enriched in NDEVs. CX3CR1 is most enriched in MDEVs, while GPR158 is the most upregulated GPCR in NDEVs. **B.** NDEVs vs. ADEVs. The orphan receptor GPRC5B shows strong enrichment in ADEVs, while GPR158 is enriched in NDEVs. **C.** MDEV vs ADEV: The chemokine receptor CX3CR1 is strongly upregulated in MDEVs. Conversely, S1PR1, GPRC5B, and GPR37L1 are highly enriched in ADEVs. The dashed-line ovals enclose the key enriched GPCRs: Orange for NDEVs, Blue for MDEVs, and Purple for ADEVs. **D.** A heatmap displaying the Log2 Fold Change (Log2 FC) of GPCRs significantly enriched in each cell type-specific EV compared to the Bulk EVs. The top-enriched receptors are GPR158 in NDEVs, CX3CR1 in MDEVs, and GPRC5B in ADEVs. GPR17 is significantly enriched in Bulk EVs. Volcano plots show gene names instead of protein names. **E.** Graphical illustration of the most enriched GPCRs by Cell type-specific EVs (created on Biorender.com).

**Figure 6.**
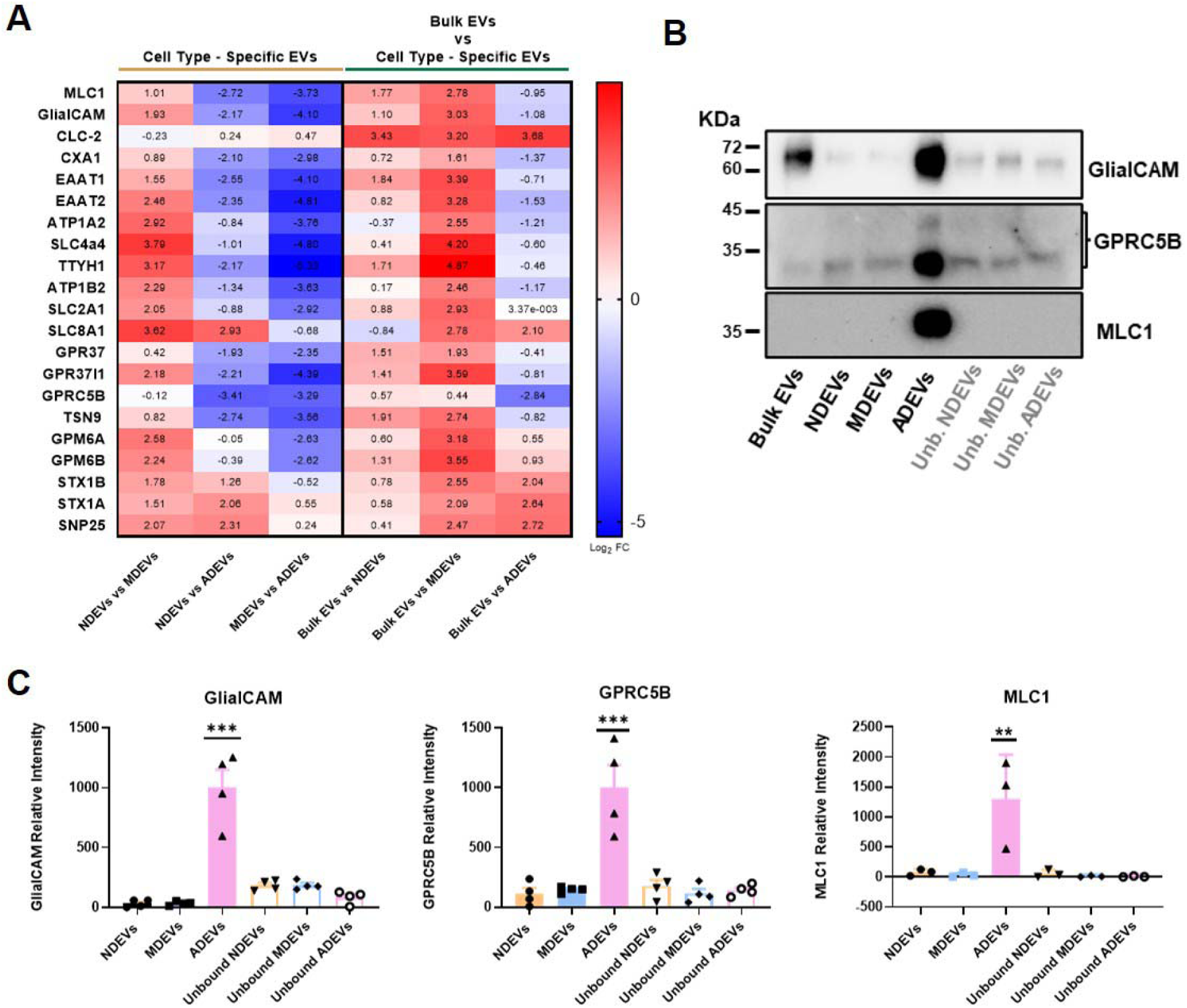
Proteomic profiling and Experimental Validation of the GlialCAM Interactome Signature in ADEVs. **A.** Heatmap illustrating the relative enrichment of GlialCAM interactome proteins in ADEVs, NDEVs, and MDEVs compared to bulk EVs and/or other cell type-specific EVs. This analysis highlights the cell type-specific distribution of astrocytic (e.g., MLC1, GlialCAM) versus neuronal (e.g., STX1A, SNAP25) markers within the interactome. **B.** Western Blot Validation of Key Astrocytic GlialCAM Interactome Proteins in ADEVs. Immunoblotting confirms the enrichment of the astrocytic markers MLC1, GlialCAM, and GPRC5B in ADEVs compared to other cell type-specific EVs (NDEVs and MDEVs). **C.** Bar graphs quantifying the relative abundance (mean ± SD) of GlialCAM, GPRC5B and MLC1, in the bound fractions (ADEVs, MDEVs, and NDEVs) versus the unbound. Data is normalized by the average intensity in ADEVs. Ordinary one-way ANOVA followed by Tukey’s multiple comparisons test, N=4 for GlialCAM and GPRC5B (*** p<0.0001); N= 3 for MLC1 (** p<0.005). Unb.: Unbound.

MDEVs were enriched in Class A chemokine receptors CX3CR1 and CCR5, purinergic receptors P2RY12 and P2RY13, and Adhesion Class receptors ADGRG1 and ADGRE1. Additional enriched receptors included the chemerin receptor CMKLR1, the complement peptide receptor C5aR1, and the orphan receptor GPR34 (Fig. 5B-C). CX3CR1 was the most significantly enriched GPCR in MDEVs compared to bulk EVs (p adj. = 7.91 × 10⁻L), NDEVs (p adj. = 5.88 × 10⁻L), and ADEVs (p adj. = 2.40 × 10⁻L; Fig. 7D-E). P2RY12, P2RY13 and GPR34 were also significantly compared to bulk EVs ( *p adj.* = 4.46 × 10 ^-2^, 5.63× 10 ^4^, and 2.29 × 10⁻³, respectively), NDEVs (*p* adj. = 8.39 × 10⁻L, 1.06× 10⁻^3^, and 6.89 × 10⁻L, respectively) and ADEVs (*p adj.* = 2.07 × 10⁻L, 4.01 × 10 ^4^, and 3.77 × 10⁻L, respectively).

ADEVs displayed selective enrichment of the Class A orphan receptor GPR37L1 and the Class C orphan receptor GPRC5B (Fig. 5B-C), together with lysophospholipid receptors S1PR1 (S1P receptor) and LPAR1 (LPA receptor), and the Class A endothelin receptor EDNRB and neurotensin receptor NTR2 (Fig. 5B-C). GPRC5B was the most significantly enriched GPCR in ADEVs compared to bulk EVs (*p* adj. = 3.10 × 10⁻L), NDEVs (*p* adj. = 1.57 × 10⁻L), and MDEVs (*p* adj. = 2.46 × 10⁻L). EDNRB was also highly enriched in ADEVs (Fig. 5D-E).

Bulk EVs, in contrast, showed exclusive upregulation of GPR17, a Class A, P2Y-like Purinergic Receptor compared to NDEVs (*p* adj. = 1.59 × 10⁻L), MDEVs (*p* adj. = 1.16 × 10⁻L), and ADEVs (*p* adj. = 6.57 × 10⁻L). Other GPCRs enriched in bulk EVs included ADGRL2 (Latrophilin 2), NTSR1, and the orphan receptor GPR107 (Fig. 5D-E).

### ADEVs Harbor a Proteomic Signature Linked to BBB Function via the GlialCAM Interactome

Astrocytes are crucial regulators of CNS homeostasis, BBB integrity, and their dysfunction is strongly associated with neurodegenerative diseases such as Alzheimer’s, Parkinson’s, and Amyotrophic Lateral Sclerosis. Given their central role in neurovascular regulation and glia–neuron communication, we aimed to identify molecular signatures within ADEVs associated with astrocyte endfeet and BBB interfaces.

To define an ADEV-specific signature, we first identified proteins consistently upregulated in ADEVs compared to bulk EVs, NDEVs and MDEVs, or only to NDEVs and ADEVs. This initial overlap analysis yielded 226 proteins (Supplementary Fig. 4). Applying a stringent threshold (FC ≥ 2 and p adj. < 0.05) reduced this set to 66 highly upregulated proteins (Supplementary Fig. 4B): 11 proteins upregulated against both Bulk EVs and other cell type-specific EVs, and 55 proteins upregulated against cell type-specific EVs only.

STRING-based protein-protein interaction (PPI) analysis and k-means clustering (k=7) revealed a major cluster related to Glial cell projection, including established ADEV markers EAAT1, EAAT2, GLUL, and AQP4 (Supplementary Fig. 4C). This cluster also contained GlialCAM-associated proteins such as GlialCAM, MLC1, GPR37L1, TTYH1, and GJA1. Other clusters included lysosphingolipid and LPA receptors (S1PR1, LPAR1, PLPP3, SPTLC1), modulator molecules of the bioactive lipid receptors activity (RGS6, ARHGEF12), postsynaptic actin remodeling (WASF3, WASF1), and adenosylhomocysteinase activity (AHCYL1, AHCYL2) (Supplementary Fig. 4C).

GPRC5B and TSPAN9, which are members of the GlialCAM interactome, were also upregulated in ADEVs but clustered outside the main Glial cell projection group. Specifically, GPRC5B clustered with Nudt3, suggesting a functional interaction based on co-expression evidence (Supplementary Fig. 4C). ATP1B2, a known astrocyte marker, was excluded from this stringent set due to FC values slightly below threshold versus Bulk EVs (FC = −1.17) and NDEVs (FC = −1.34).

Analysis of GlialCAM interactome membership confirmed supported cell type-specificity in protein subtype-specific enrichment: ADEVs were enriched for adhesion and transport proteins (e.g., EAAT1-2, MLC1), while NDEVs showed enrichment for synaptic junctions within the same interactome (e.g., STX1A, STX1B, SNAP25; FC vs ADEVs = 2.06, 1.26 and 2.31, respectively). GPM6A and GPM6B were similarly abundant in both ADEVs and NDEVs (Fig. 6A).

Western blot validated the significant enrichment of the three key GlialCAM interactome components in ADEVs: MLC1, GlialCAM and GPRC5B (Fig. 6B-C). These findings suggest that ADEVs act as carriers of molecular machinery essential for BBB support. Specifically, MLC1 and GlialCAM maintain astrocyte endfeet structure and BBB integrity, while GPRC5B enrichment indicates potential roles in receptor-mediated communication pathways for brain homeostasis.

## DISCUSSION

EVs secreted by brain cells represent a key interface for neuro-glial communication and a promising source of biomarkers for CNS disorders. By comprehensively characterizing neuronal-, microglial-, and astrocyte-derived EVs (NDEVs, MDEVs, and ADEVs) from mouse brain tissue, we delineate a proteomic landscape that captures the molecular diversity of brain EVs and provides insight into astrocytic pathways of BBB regulation and neurodegeneration.

Each EV subtype exhibited molecular fingerprints consistent with parental cell functions. NDEVs were enriched in synaptic and vesicle trafficking proteins such as APP, SNAP25, L1CAM, NCAM1, and BDNF, reflecting roles in cell adhesion, neurotransmission, and synaptic plasticity. Although bulk EVs were highly similar to NDEVs, our data indicate that multiple neuronal subtypes (both excitatory and inhibitory) from diverse brain regions contribute to the NDEV pool. MDEVs contained immune and phagocytic markers including bona fide microglial markers such as CX3CR1, P2RY12/13 and CCR5, consistent with their functions in neuroimmune signaling and debris clearance. ADEVs were characterized by astrocytic transporters and endfoot-associated proteins such as EAAT1/2, AQP4, and GJA1, highlighting their involvement in neurotransmitter recycling, metabolic homeostasis, and BBB maintenance. Furthermore, our data showed that, even though bulk EVs were similar to NDEVs, most neuronal types from many areas were highly represented in our NDEVs. To explain this similarity, we can think that neurons release the vast majority of EVs in the brain, challenging to reach saturating conditions in the immunocapture; alternatively, NDEVs are highly heterogeneous and express distinct surface markers. In the first case, this will explain why bulk EVs were not enriched with oligodendrocytic EVs, and in the latter case, this will justify the apparent controversy regarding the use of different markers to pull down NDEVs in the literature. These findings align with previous reports demonstrating distinct proteomic profiles of neuron-, microglia-, and astrocyte-derived EVs in hiPSC-derived brain cells ^15^ and human brain tissue where neuronal (SNAP25, GRM7, GPR158) and astroglial markers (AQP4, EAAT1/GLAST) have been detected in total EV preparations ^35,36^ and linked to neurodegenerative pathology. Comparison with the ExoCarta Top 100 small EV proteins confirmed shared EV architecture among subtypes, yet revealed cell-type-dependent distribution: flotillins (FLOT1, FLOT2) were enriched in NDEVs and ADEVs, suggesting flotillin-positive membrane microdomains as EV origins ^37,38^, whereas MDEVs showed higher levels of ALIX and TSG101, indicating a stronger reliance on ESCRT-dependent endosomal biogenesis^39^. These differences support the existence of multiple EV biogenesis routes across brain cell types that shape their molecular cargo and signaling functions, and should be taken into consideration when selecting EV markers for characterization and validation of brain cell-specific EVs.

A striking observation was the broad representation of G protein–coupled receptors (GPCRs) across EVs, underscoring the importance of receptor-mediated signaling in EV biology. NDEVs were enriched in class C receptors including the metabotropic glutamate receptor family and GPR158, linked to synaptic signaling and cognitive regulation ^40–42^; MDEVs contained chemokine and purinergic GPCRs such as CX3CR1, GPR34, P2RY12, and P2RY13, consistent with their immune-sensing roles ^43–47^; and ADEVs exhibited selective enrichment of the astroglial orphan receptors GPR37L1 and GPRC5B, both implicated in glial stress responses and neuroprotection ^22,48^. The presence of GPCRs in EVs suggests that they may act as carriers of functional receptors, modulating intercellular communication and potentially participating in feedback regulation of signaling pathways ^33,34,49^. Importantly, EV-associated GPCRs (EV-GPCRs) have been linked to various pathological contexts ^33,49–51^, including Alzheimer’s and Tau pathologies ^52,53^, positioning them as promising biomarker candidates for neurodegenerative disorders.

Within ADEVs, we identified a coherent network of GlialCAM-associated proteins, including MLC1, GlialCAM, ATP1B2, EAAT1/2, TTYH1, GJA1, GPRC5B and GPR37L1, representing a molecular signature of astrocytic endfeet and their interactions with endothelial and pericyte cells at the BBB ^22,54^. Supporting our results, GlialCAM and GPRC5B were also found in ADEVs from in vitro human IPSCs cultures ^15^. Furthermore, this observation converges with recent studies showing that astroglial exosomal GlialCAM mediates axonal growth and neuroprotection, while inflammatory cytokines reduce GlialCAM-positive EV release in mouse models of amyotrophic lateral sclerosis ^55^. The detection of GPRC5B in ADEVs aligns with its previously reported exosomal sorting and intercellular transfer ^49,56^, suggesting that GPCRs within the GlialCAM interactome may travel in EVs and influence neurovascular signaling. The overlap between ADEVs cargo and the GlialCAM/MLC1 interactome also links EV composition to ion and water homeostasis pathways implicated in megalencephalic leukoencephalopathy with subcortical Cysts (MLC), supporting the hypothesis that ADEVs could reflect functional states of astrocytic endfeet relevant to BBB integrity and white-matter homeostasis. Notably, GPRC5B and GPR37L1, validated components of the GlialCAM interactome, modulate MLC1 and GlialCAM levels, and GPRC5B has been associated with MLC genetics ^22^, reinforcing its translational interest. Several members of this network, including EAAT1 ^15,16^, EAAT2, and ATP1B2 ^17^, are cell-surface proteins already used for ADEV isolation, highlighting their immuno-targeting potential. In contrast, although GlialCAM has been detected in EVs, it has never been used directly as an immunoprecipitation target ^55,57^. MLC1, while less studied in the EV field, is listed in EV proteome databases ^58^ and localizes to membrane, endosomal, and multivesicular body (MVB) compartments in astrocytes ^59^, which makes it a candidate for future validation as an EV-associated protein or immunocapture target.

Integrating the ADEVs proteome with GPCR and canonical EV marker profiles further reveals a complex interplay between vesicle biogenesis pathways, signaling functions, and neurovascular specialization. Additional members of the GlialCAM-associated GPCR family, such as GPR37L1, have been identified as EV-associated proteins in EV-enriched populations spontaneously released from mouse and human brain tissue ^60^, and GPRC5B has similarly been detected in EVs across multiple studies ^15,49,56^. Our results, together with these previously reported associations of GlialCAM-linked GPCRs with EVs, further support their conserved localization within astrocyte-derived EV populations.

While our dataset provides a robust reference map for brain cell–specific EVs, some limitations should be acknowledged. The use of young adult brain tissue may underestimate age-related or pathology-specific alterations. Moreover, regional and subtype heterogeneity within astrocyte and microglial populations could contribute to undetected EV diversity. We also note that GPM6A, previously reported as a neuronal EV marker ^36^, was enriched in both NDEVs and ADEVs, potentially reflecting overlapping molecular characteristics between these vesicle populations. Future studies incorporating spatial proteomics, single-vesicle analyses, and disease models will be critical to define how ADEVs composition changes in response to BBB breakdown and neurodegenerative progression.

In summary, our comparative proteomic profiling establishes a detailed atlas of neuronal-, microglial-, and astrocytic-derived EVs, offering a valuable resource for target selection and cargo characterization. Furthermore, we report that ADEVs exhibit a distinctive molecular signature enriched in BBB-associated proteins and GPCRs, highlighting astrocytic specialization in neurovascular and neuroprotective signaling. Together, these findings provide a molecular foundation for exploring cell type-specific biomarkers in EVs, and for understanding the vesicular mechanisms linking astrocyte function to BBB integrity and CNS disorders.

## ACKNOWLEDGEMENTS

We are grateful to the Microscopy Unit at ISABIAL, with the technical support of Guizlane Ennatiji, Andrea Ibáñez, and Andrea Poveda, and the Proteomics facility of SCSIE University of Valencia for their valuable help with EM and proteomics analysis, respectively. We would like to acknowledge José Amable Bernabé’s contribution for particle size distribution and concentration measurements at the Barcelona Materials Science Institute (Unit 6, ICTS “NANBIOSIS”, part of CIBER-BBN).

## ETHICS APPROVAL STATEMENT

All mouse procedures were approved by the Comité de Ética de la Investigación, Universidad de Alicante, and authorized by the Dirección General de Producción Agrícola y Ganadera, Generalitat Valenciana, according to European, national and regional laws.

