## Supplementary Figures for "Cell type-specific Extracellular Vesicles in Mouse Brain: Proteomic Signatures Highlight Astrocytic GlialCAM Network and GPCR Enrichment"

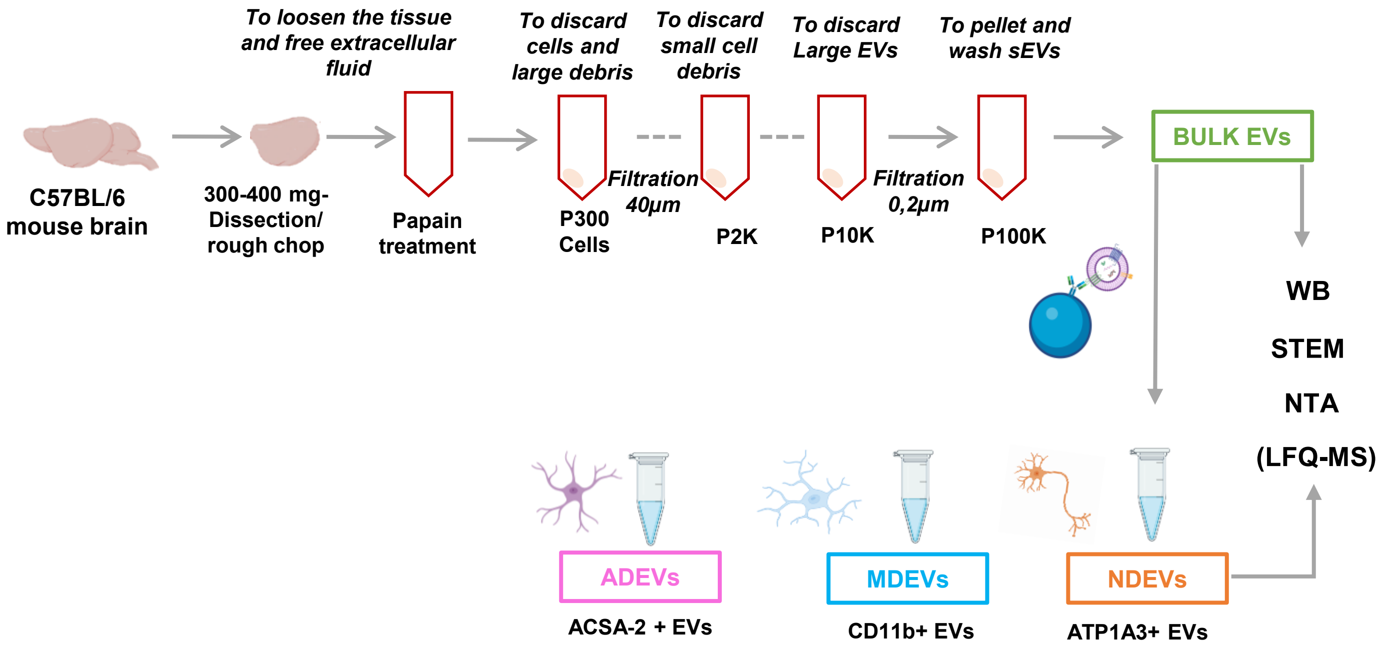


**Supplementary Figure 1. Workflow of Bulk and Cell Type–Specific Extracellular Vesicle Isolation.** Schematic overview of the experimental workflow used to isolate bulk EVs and immunocaptured cell type–specific EVs from mouse brain for proteomic characterization.

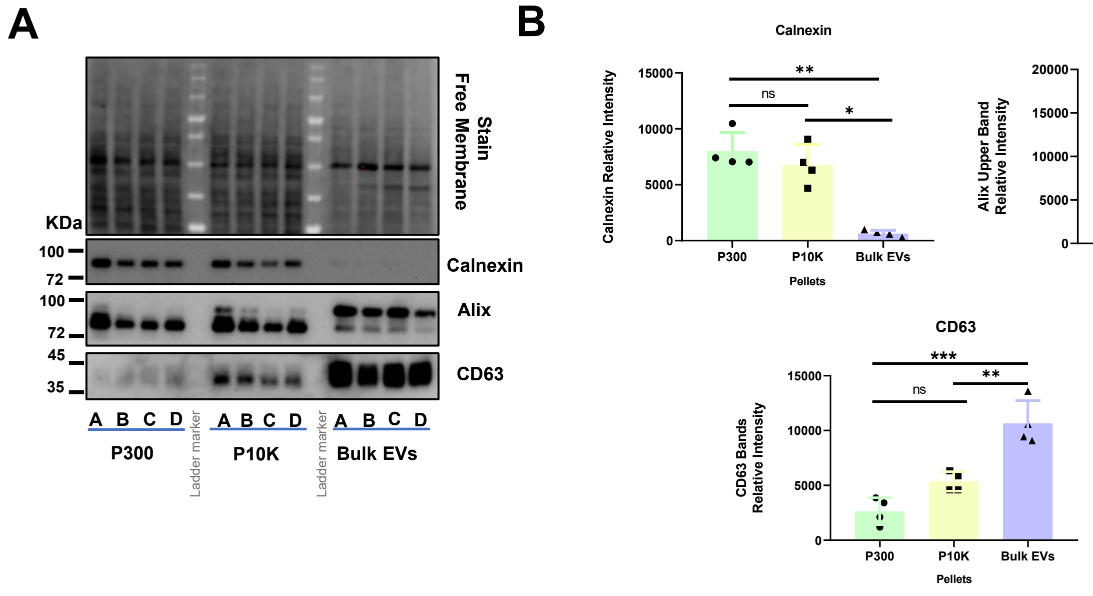


**Supplementary Figure 2. Validation of Bulk Extracellular Vesicle Isolation**. **A.** Representative Western blot showing enrichment of the EV markers Alix and CD63 in the Bulk EVs fraction compared to P10K and P300 fractions, accompanied by a marked reduction of the endoplasmic reticulum marker Calnexin (N = 4). **B**. Quantification of Western blot bands shows significantly lower Calnexin levels in P100K compared to P300 and P10K with no difference between P300 and P10K. Alix expression was significantly higher in Bulk EVs relative to P300 and P10K , with no difference between P300 and P10K. CD63 was enriched in Bulk EVs compared to both P300 and P10K, while P10K showed higher CD63 levels than P300. These data confirm the enrichment of small EV-associated markers in the Bulk EVs and the progressive depletion of cellular contaminants with increasing centrifugation force. Ordinary one-way ANOVA followed by Tukey's multiple comparisons test were used for the Western blot data. Data is shown as mean ± SD (Standard Deviation), * $P < 0.05$, ** $p<0.01$, *** $p < 0.001$. P300= Pellet 300 × g; P2K: Pellet 2000 × g; P10K = Pellet 10,000 × g. P100K or Bulk EVs= Pellet 100,000 × g. Neuronal- (NDEVs), Microglial- (MDEVs), and Astroglial-Derived EVs (ADEVs).

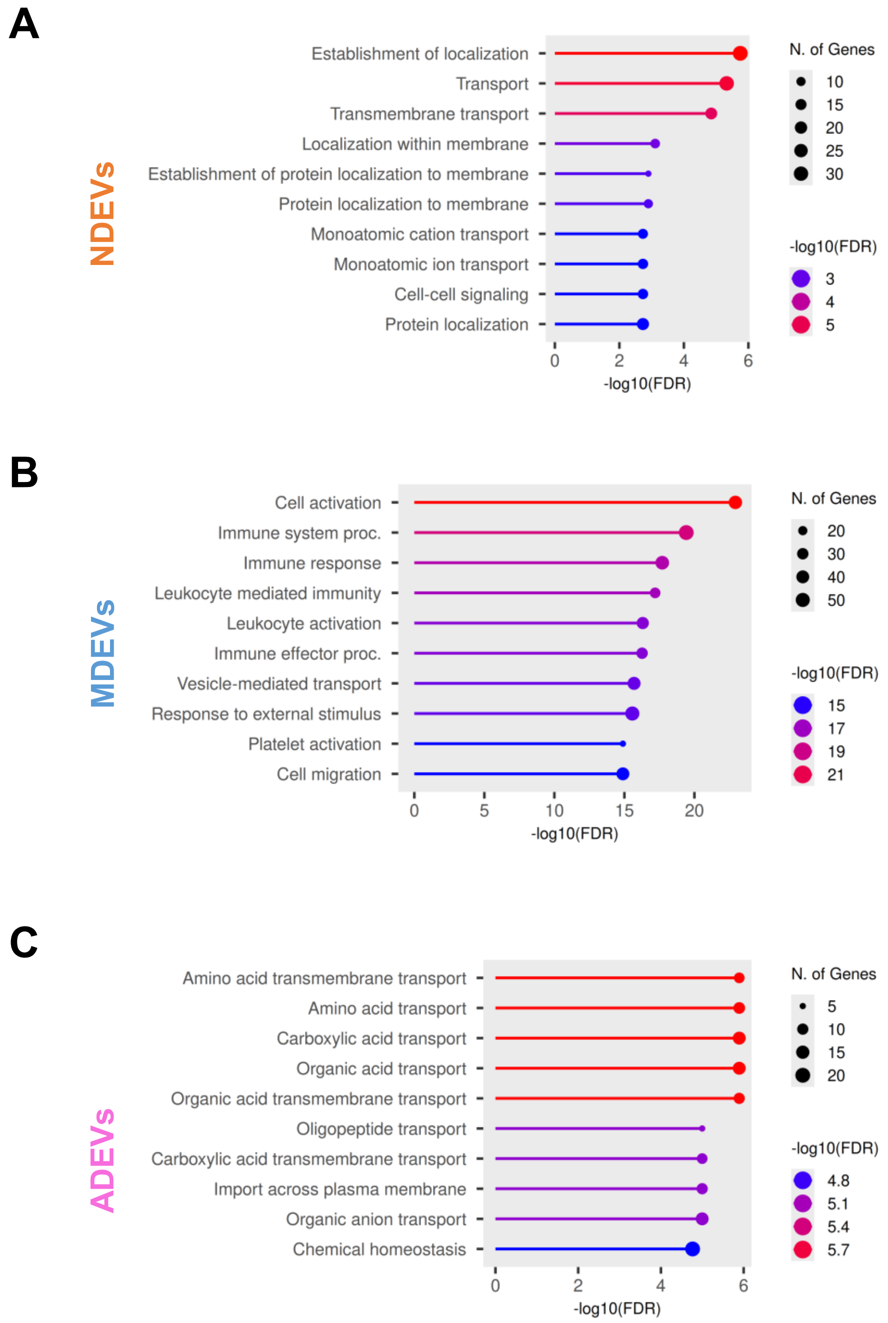


**Supplementary Figure 3. GO Biological Process of Cell-Type-Specific EV Subtypes. A.** GO Biological Process enrichment analysis of the 58 NDEV-upregulated proteins, highlighting pathways related to establishment of localization, transmembrane and ion transport, and cell–cell signaling. **B.** GO Biological Process analysis of the 119 MDEV-upregulated proteins, demonstrating enrichment in cell activation, immune system processes, immune response, and vesicle-mediated transport. **C.** GO Biological Process enrichment of the 93 ADEV-associated proteins, showing pathways related to amino acid transmembrane transport, carboxylic and organic acid transport, and chemical homeostasis.

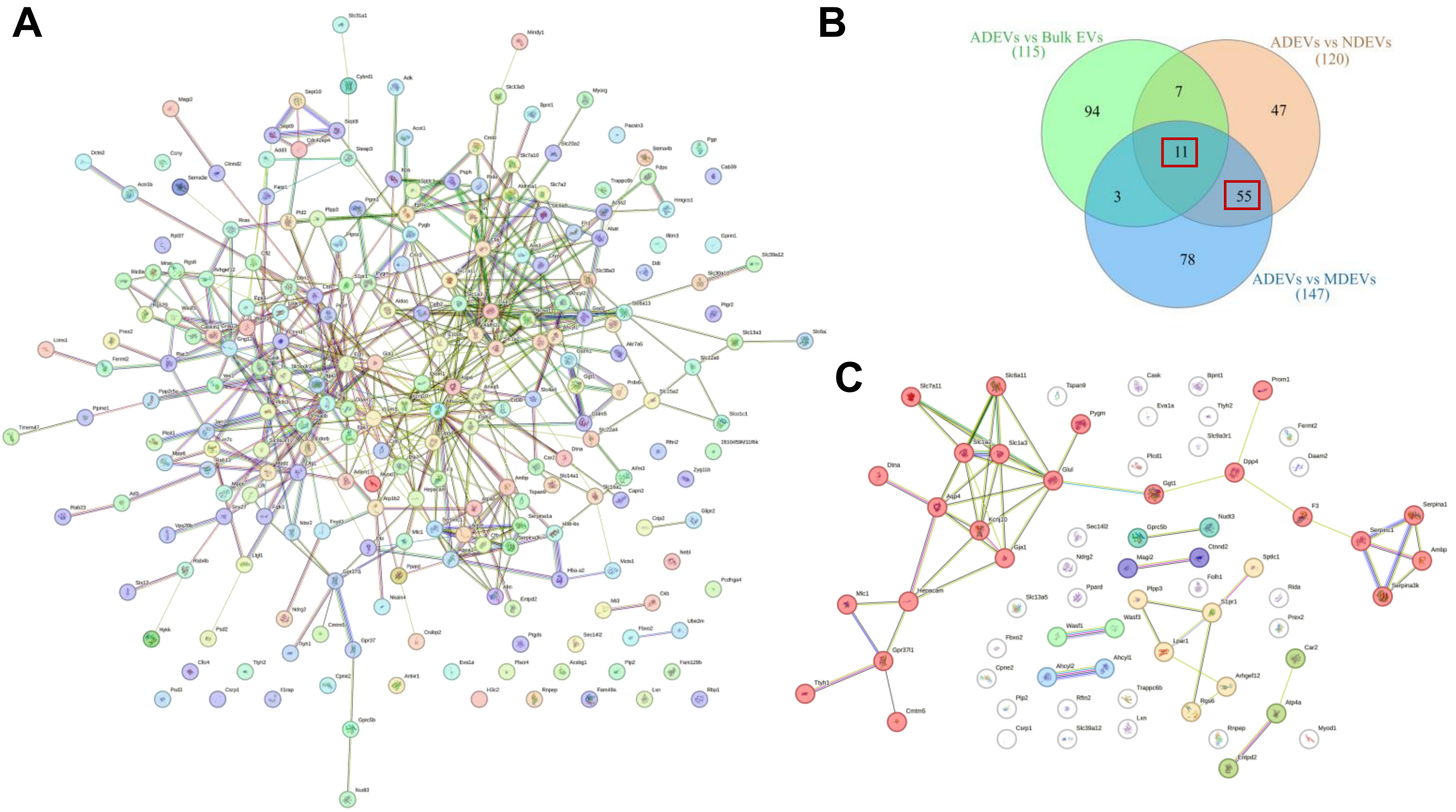


**Supplementary Figure 4. Detailed Proteomic Network Analysis of Proteins Upregulated in ADEVs. A.** Protein-Protein Interaction (PPI) Network including the full set of **226 proteins** identified as consistently upregulated in ADEVs (FC ≥ 1 p adj. < 0.05). This set includes the 93 proteins enriched in ADEVs versus Bulk EVs and Cell-Type Specific EVs, plus the 133 proteins enriched only compared to the Cell-Type Specific EVs. **B.** Venn Diagram illustrating the overlap of proteins meeting the stringent functional threshold criteria (FC ≥ 2 p adj. < 0.05) when comparing ADEVs to Bulk EVs and Cell-Type Specific EVs. The diagram highlights the resultant 66 highly upregulated ADEV proteins. **C.** Protein-Protein Interaction (PPI) Network visualizing the interaction map of the 66 highly upregulated ADEVs proteins, confirming their unique molecular profile relative to all control groups.
